# Groundwater and phenology data reveal vulnerability of riparian trees to drought

**DOI:** 10.1101/2025.05.23.654357

**Authors:** Rose M. Mohammadi, Todd E. Dawson, Claire R. Tiedeman, John C. Stella, Albert Ruhí

## Abstract

The increasing frequency and magnitude of climatic extremes are altering water availability in dryland ecosystems globally. However, riparian vulnerability to hydroclimate whiplash remains poorly understood. Here, we examined how riparian trees respond to groundwater fluctuations and drought through their water use patterns and phenology. To this end, we combined time-series analysis of long-term, high-frequency groundwater monitoring and satellite imagery from a drought-prone and relatively pristine watershed in California (Chalone Creek, Pinnacles National Park). We found that trees by intermittent river reaches displayed consistent but depth-limited groundwater reliance, while those at perennial reaches primarily relied on groundwater during the dry season. Machine-learning models revealed that at intermittent sites groundwater depth predominantly controlled vegetation greenness, represented by Normalized Difference Vegetation Index (NDVI). In contrast, variation in photoperiod length dominated at perennial sites where water was more reliably available. During the severe 2020-2022 drought, all species experienced reduced greenness, but phenological responses differed by flow regime. While the start of season was delayed across all sites, trees at intermittent reaches exhibited substantially earlier end of season during drought, resulting in growing seasons shortened by as much as 28 days. These phenological shifts vastly exceed those documented across aridity classifications in global datasets from satellite observations, ground-based monitoring networks, and experimental precipitation manipulations. Although riparian trees in drylands have been shaped by exposure to drought over evolutionary timescales, our findings challenge the prevailing assumption that ecosystems regularly exposed to hydrological are more resilient to drought. Instead, we show that trees in intermittent systems may be operating close to critical groundwater thresholds, rendering them particularly vulnerable to increasingly long and severe droughts.

## Introduction

Increasing climate variability, including more frequent and extreme droughts punctuated by exceptionally wet years, is reshaping water availability in dryland ecosystems globally. In California, this hydroclimate whiplash is driven by intensifying atmospheric river activity and warming temperatures, resulting in precipitation extremes that are unprecedented in the past 600 years (Dettinger, 2011; Swain et al., 2025). This volatility poses a serious risk to dryland riparian zones, which, despite covering a small percentage of land area, contribute a disproportionately large fraction of landscape-scale primary productivity and evapotranspiration (Doody et al., 2023). Riparian corridors also are biodiversity hotspots, serving as thermal and moisture refuges that support different assemblages than those found in aquatic or terrestrial environments as well as with many rare and endemic species (Sabo et al., 2005; Cartwright et al., 2020). However, these ecosystems are threatened by changes in water availability, including alterations of natural flow regimes and declines in groundwater levels (Stromberg et al., 1996; Stromberg, 2001; Williams & Cooper, 2005). Despite their ecological importance, the vulnerability of dryland riparian zones to interannual variation in hydroclimate remains poorly understood.

Dryland riparian vegetation is adapted to localized and fluctuating water availability, including periods of low and high flow (Lytle & Poff, 2004). Different species cope with water deficits differently, along a continuum from anisohydric to isohydric responses (McDowell et al., 2008; Hultine et al., 2020; McDowell et al., 2022). Anisohydric species seek to tolerate drought by keeping stomata open to maintain a positive carbon balance, but that strategy comes at the risk of low leaf water potentials and thus potential hydraulic failure. In contrast, isohydric species close their stomata or shed leaves to keep leaf water potentials relatively constant, avoiding short-term cavitation-induced mortality but risking long-term carbon starvation instead (Rood et al., 2003; McDowell et al., 2008; Zunzunegui et al., 2011). These strategies may not always suffice in light of intensifying droughts, which often push riparian ecosystems beyond resilience thresholds (e.g., branch abscission, die-backs) (Stella et al., 2013; Stella & Bendix, 2019). Phenological shifts such as changes in the timing of leaf out, flowering, fruiting, and senescence can help reduce drought stress (Menzel et al., 2006; Wada et al., 2014; Piao et al., 2019), and increasing evidence suggests that droughts can prolong senescence in dryland riparian ecosystems (Kibler et al., 2021; Warter et al., 2021). Because climate and non-climate stressors often co-occur, understanding how climatic extremes induce phenological change remains a challenge.

Groundwater, or water stored in the saturated zone, is a critical resource in dryland riparian ecosystems. In intermittent streams that stop flowing or completely dry up for part of the year, shallow groundwater can support critical refugia for aquatic food webs, including invertebrates, fish, and amphibians that require long hydroperiods (Bogan et al., 2017; Palmer & Ruhí 2019). In terrestrial ecosystems, groundwater sustains phreatophytes, plants that rely on groundwater (Orellana et al., 2012; Hultine et al., 2020). During dry periods, groundwater can become decoupled from vadose zone soil moisture, providing an alternative water source and allowing these deeply rooted plants to escape the effects of dry soil moisture conditions (Gou & Miller, 2014; Quichimbo et al., 2020). Further, temporal variation in the availability of the different water sources can influence the phenology of key life history events. For example, willows and cottonwoods (Salicaceae) synchronize propagule dispersal with peak or receding spring flows (Stella et al., 2006), and floods further promote their dispersal via root and shoot regeneration (Karrenberg et al., 2002). While temperature is generally considered the primary control of plant phenology in temperate environments (Cleland et al., 2007; Piao et al., 2019), in water-limited dryland ecosystems, moisture availability can control vegetation growth and annual peaks and troughs of greenness (Rodriguez-Iturbe, 2000; D’Odorico et al., 2007; Warter et al., 2023). Therefore, drought-induced changes in moisture availability arising from variation in groundwater depth and surface streamflow could greatly alter riparian tree phenology along river corridors.

Understanding how riparian vegetation interacts with fluctuations in groundwater across intermittent river networks requires capturing both fine-scale temporal variation and spatial context. Intermittent watersheds are highly dynamic and spatially heterogeneous (Datry et al. 2016), with narrow stream corridors that experience rapid hydrologic shifts. High-frequency groundwater depth measurements can offer key insights into plant-water interactions, particularly when paired with remote sensing tools that track vegetation change over time (Elmore et al., 2006). Integrating time-frequency methods like wavelet transforms can reveal hydrologic signatures that would be missed using standard time-domain approaches (Yu et al., 2015; Ruhí et al., 2018). Combining these techniques with machine learning may allow nuanced understanding of species-specific water use strategies, and how different plants respond to water stress (Ruehr et al., 2023; Qiao et al., 2024). In this study, we applied a novel combination of time-series and machine-learning methods to understand vegetation response across temporal scales, and to identify the environmental drivers of phenological change.

California’s hydroclimate over the last decade provides a unique opportunity to examine how riparian vegetation responds to rapid swings in water availability. The state has recently experienced record-setting drought years—water years 2021 and 2022 were the second and third driest on record in California (California Department of Water Resources, 2021; California Department of Water Resources, 2022)—and exceptionally wet years (2019, 2023, 2024), creating ideal conditions to study how riparian trees change their water use strategies and phenology in response to shifting moisture availability. Our study focuses on an unmodified, highly instrumented river network, Chalone Creek (Pinnacles National Park, California), which has a gradient of streamflow permanence and varying groundwater regimes. We explored three questions: (1) How do the timing and magnitude of groundwater use by riparian trees vary with flow regime and season across the stream network? (2) How does variation in depth to groundwater influence riparian vegetation phenology in intermittent and perennial stream reaches, and how does its importance compare to other environmental drivers of phenology and greenness? (3) How sensitive are water-riparian vegetation interactions to drought, and how does this sensitivity compare to other ecosystems globally?

We hypothesized that riparian trees near perennial stream reaches primarily rely on groundwater during the dry summer but use soil moisture recharged by precipitation and groundwater discharge in the wet spring. In contrast, trees near intermittent stream reaches, where the vadose zone is less consistently recharged due to deeper and more variable groundwater levels, may depend on deeper phreatic water sources year-round (Snyder & Williams, 2000; Sargeant & Singer, 2016; Gou & Miller, 2014; Flanagan et al., 2019; Williams et al., 2024). Consequently, we expected depth to groundwater and precipitation to exert the strongest influence on vegetation canopy greenness (Huntington et al., 2016; Rohde et al., 2021). We also predicted that supraseasonal drought would shorten the growing season by accelerating leaf senescence, with this effect being more pronounced at intermittent than at perennial sites, due to more limited groundwater access in the latter (Rood et al., 2000). Finally, we hypothesized that dryland riparian ecosystems may be more sensitive to drought, due to their strong reliance on variable groundwater, than more humid ecosystems in other climates with more stable moisture availability. Thus, we expected that supraseasonal drought would induce strong phenological shifts (e.g., earlier leaf senescence) in dryland riparian trees, revealing the vulnerability of these critical ecosystems to the ongoing intensification of the hydrologic cycle.

## Methods

### Study Site

Chalone Creek, Pinnacles National Park, California is an intermittent watershed where about ∼90% of the stream network naturally dries for part of the year. Stream reaches along the network span a broad range of hydroperiods (from ∼15% to 100% flow permanence), and groundwater discharges from several springs throughout the park, which in the dry season are the sole source of flow in the few perennial reaches. The mainstem of Chalone Creek is a losing stream—the creek tends to recharge the groundwater system during wet-season periods when precipitation is minor or absent—while its main tributary Sandy Creek is a gaining stream, with groundwater discharging to surface water flow. In both alluvial areas, groundwater flows southerly, following the creek drainages. The aquifer is recharged by precipitation and inflow from alluvium upstream (Scheiderich et al., 2022).

The watershed has a Mediterranean climate, with hot, dry summers and cool, moderately wet winters, and most precipitation falling October to May. We delineate the wet and dry seasons using precipitation data whereby the wet season begins when accumulated precipitation in the water year exceeds 20 mm and ends with the last day of precipitation before the summer (June 1). The dry season is the period between wet seasons. Average annual precipitation was 414 mm from 1937 to 2023 (Western Regional Climate Center 2024a, Western Regional Climate Center 2024b). The study period includes both wet and dry years compared to the long-term average: water years 2020 (300mm), 2021 (223mm), and 2022 (225mm) received below-average rainfall, consistent with water years 2021 and 2022 being the second and third driest on record in California (California Department of Water Resources, 2021; California Department of Water Resources, 2022). In turn, water years 2019 (446mm), 2023 (581mm), and 2024 (426mm) received above-average rainfall.

### Groundwater data collection and processing

Depth to groundwater data (DTW, m) were collected from six monitoring wells at 15-minute intervals from January 2019 to November 2024 (Figure 1A), three along the mainstem of Chalone Creek (CHA-2, CHA-3, and CHA-4) and three on the Sandy Creek tributary (SC-2, SC-4, and SC-5). All wells were located on intermittent reaches except SC-5. Groundwater levels were measured at CHA-2, CHA-3, CHA-4, SC-4, and SC-5 using Rugged TROLL 100 data loggers, which include an absolute pressure transducer (In-Situ Inc). Barometric pressure was recorded at a 15-minute frequency using an In-Situ Baro TROLL about 0.3 m below land surface in well CHA-2. Groundwater pressure data were processed to remove barometric effects and converted to depth measurements. In SC-2, groundwater data were collected using a vented pressure transducer (Process Measurement & Controls Inc. Miniature Submersible Depth / Level Transducer model MTM 3211) connected to a Campbell Scientific CR10x datalogger at land surface. Field visits to download data and manually check groundwater measurements for accuracy using Solinst 102M Mini Water Level Meter were conducted biannually beginning in 2018. Further information on well specifications can be found in Scheiderich et al. (2022).

**Figure 1.**
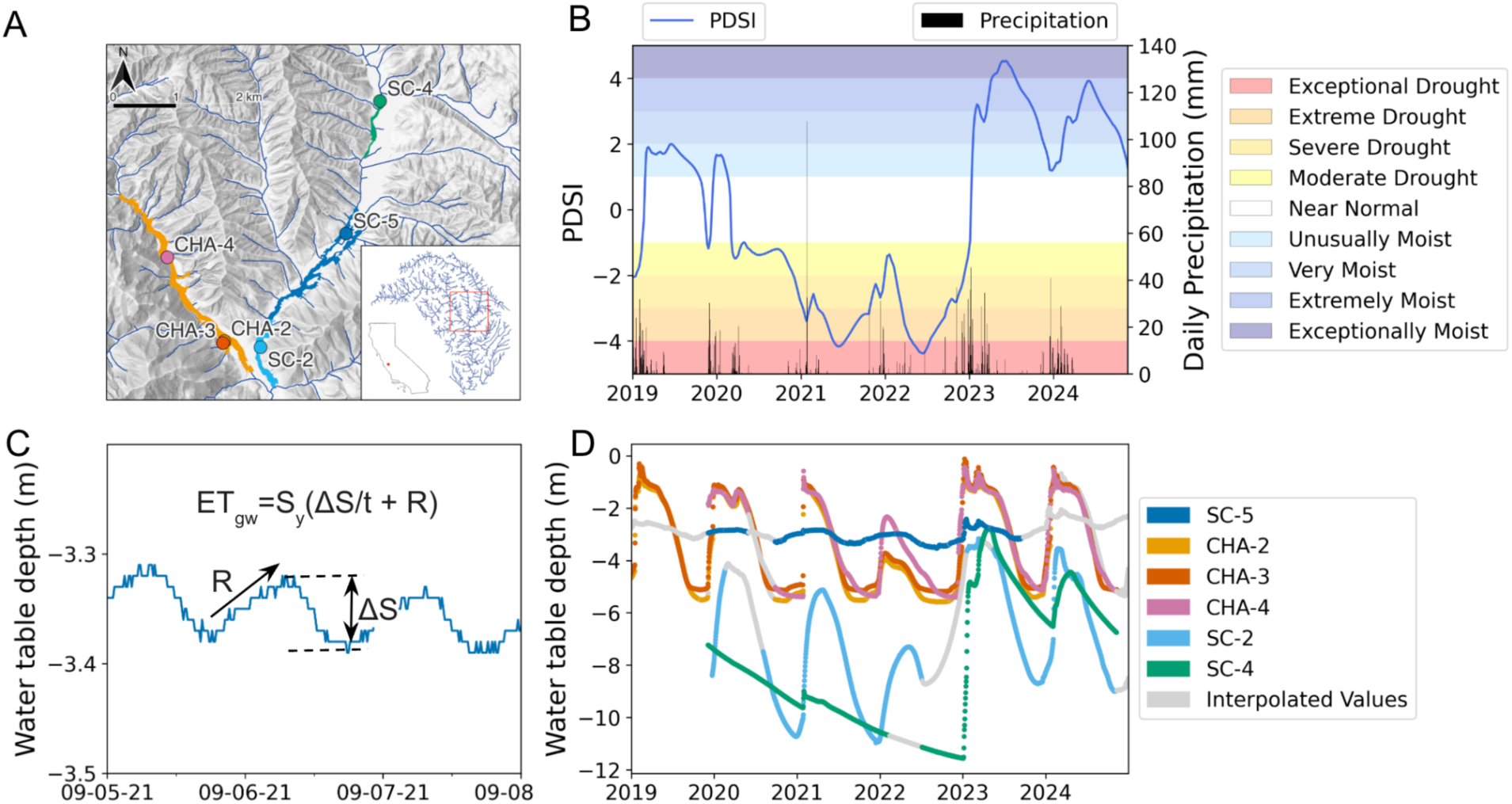
Study site and time series of long-term, high-frequency groundwater data. (A) Map of the 6 monitoring wells (circles) on Chalone Creek (Pinnacles National Park, California) and their corresponding riparian areas of influence (delineated by color). CHA wells are in the Chalone alluvial area, an intermittent, losing reach. SC wells are in the Sandy Creek alluvial area, a gaining reach where only SC-5 is perennial. (B) 5-day Palmer drought severity index (PDSI) and daily precipitation (mm). Color bands indicate corresponding qualitative drought classifications from the National Weather Service. (C) 3 days in September 2021 showing diel water table fluctuations and implementation of the White method to determine the amount of evapotranspiration coming from groundwater (ET_gw_). The daily change in storage (S) and the recovery rate (R) are used to calculate ET_gw_ for a 24-hour period (t). (D) Water table depth (m) time series at the 6 monitoring wells. All wells except SC-4 exhibited diel water table fluctuations. Each year, groundwater declined to its deepest point at the end of the dry season and rises with the onset of precipitation. The 2020-22 period captured a historic drought, with water years 2021 and 2022 being the second and third driest on record in California (California Department of Water Resources, 2021; California Department of Water Resources, 2022).

When there were gaps in the groundwater time series (average completeness was 82.39%), we used autoregressive integrated moving average models (ARIMA) and a Kalman filter to impute missing values in a way that respected the dominant periodicities, trends, and noise in the time series (Comte et al., 2021) using the ‘forecast’ package (Hyndman et al., 2024) in R (R Core Team, 2021). Across the five wells, the root mean square error (RMSE) ranged from 0.018 to 0.208 m, indicating strong agreement between modeled and observed groundwater levels. This process delivered five full years of 15-min data (January 1, 2020 to December 31, 2024).

### Meteorological data collection and processing

Daily mean air temperature (Temp, °C), daily precipitation (Precip, mm), and mean vapor pressure deficit (VPD, kPa) were collected by a weather station in the park (Western Regional Climate Center, 2024b). Accumulated precipitation over the preceding month (Precip_1m, mm) was calculated from the precipitation data. Daily photoperiod (Photo, hours) was retrieved using the U.S. Naval Observatory’s “Duration of Daylight/Darkness Table for One Year” (U.S. Naval Observatory, 2024). Palmer Drought Severity Index (PDSI, unitless)—a measure of dryness used to monitor droughts that ranges from −10 (dry) to 10 (wet)—were sourced from the Gridded Surface Meteorological (gridMET) dataset retrieved from Climate Engine (Abatzoglou, 2013) (Figure 1B). PDSI data were reported at five-day intervals and were upsampled to a daily frequency via a Kalman filter.

### Vegetation data and satellite image processing

We compiled and processed Analysis-Ready PlanetScope imagery, a 3-meter orthorectified surface reflectance product imagery, using Planet’s API (Planet Labs PBC, 2018). This imagery’s spectral bands and spectral response functions are equivalent to the blue (B2), green (B3), red (B4), and narrow NIR (B8a) bands of Sentinel-2. All images acquired during the 6-year study period that have less than 90% cloud cover were included in the study (n=746). Images were masked with Planet’s usable data mask.

We used a random forest classification model, a machine learning algorithm composed of an ensemble of decision trees, to identify deciduous vegetation canopy pixels by species and mask out evergreens, scrub, and bare ground (Breiman, 2001). We used high-resolution Google Earth imagery (from February 2021, March 2022, and July 2023) to manually draw 350 polygons (totaling 55,736 pixels) representing tree crowns, scrub and non-woody vegetation, buildings, roads, and bare ground. The polygons were classified using visual interpretation as well as 100 GPS coordinates taken next to trees along the riparian corridors of the park. The resulting pixels were randomly divided into a training (75%) and testing set (25%) to train and validate the model. Bands 1 (coastal blue, 443nm), 2 (blue, 490nm), 3 (green I, 531nm), 4 (green, 565nm), 5 (yellow, 610nm), 6 (red, 665nm), 7 (red edge, 705 nm), and 8 (NIR, 865 nm) from 8-band PlanetScope imagery acquired in January, March, and November of 2024 were used to train the model. The accuracy of each model was validated with the testing dataset. We ran the random forest model with the ‘randomForest’ package (Liaw & Wiener, 2002). The resulting model was used to classify pixels belonging to the dominant deciduous riparian tree species in the watershed: *Salix laevigata* or *Salix lasiolepis* (red and arroyo willow, “willow”), *Populus fremontii* (Fremont’s cottonwood, “cottonwood”), and *Quercus lobata* (valley oak, “valley oak”). All other pixels not classified as one of the three target species, such as the evergreen *Quercus agrifolia* (coast live oak) and *Pinus sabiniana* (gray pine), were masked out and excluded from analysis.

We calculated Normalized Difference Vegetation Index (NDVI), a metric of vegetation “greenness,” for each deciduous pixel in each image. NDVI (unitless) is calculated using the near-infrared (NIR) and red (RED) bands:

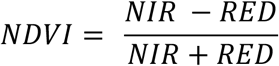

NDVI ranges from −1 to 1, with values closer to 1 indicating vegetation with a higher density of green leaves, positive small values (between 0 and 0.20) indicating no vegetation, and negative values denoting the presence of surface water. The daily average NDVI for each tree species within the 4 study reaches (Figure 1A) was computed. The resulting 12 time series were smoothed using a weighted HANTS (Harmonic Analysis of NDVI Time Series) smoother in the ‘phenofit’ R package (Kong et al., 2022). The weighted HANTS algorithm, based on the Fourier transform, reduces noise in NDVI time series by applying a moving support domain to assign weights, thereby determining frequency more easily, resulting in a time series that fits the actual vegetation growth profile (Verhoef, 1996; Yang et al., 2015).

After applying the weighted HANTS smoother for rough fitting, growing season division was performed with the ‘phenofit’ package using the threshold method. Where the original vegetation time series is *f*(*t*), we set the threshold *f*(*t*)*_ratio_* to 0.5, and define it as the normalized vegetation time series in the range of [0, 1]:

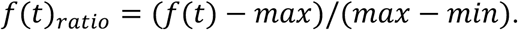

The threshold method defines the start of the growing season (SOS) and end of the growing season (EOS) by the first day when the vegetation index value exceeds and falls below the threshold— here, 50% of the seasonal amplitude—respectively. The growing season length (GSL) is the number of days between the SOS and EOS.

### Testing our hypotheses

We analyzed DTW to test our first hypothesis that riparian trees near perennial streams rely more heavily on groundwater in the dry summer, while trees near intermittent streams rely on groundwater year-round. Specifically, we sought to determine the timing and magnitude of groundwater use by vegetation using wavelet analyses. Wavelet analysis is a time-series method that decomposes signals to extract dominant frequencies, amplitudes, and phases in the data, as well as any changes in these properties over time (Mallat, 1989; Torrence & Compo, 1998). This technique has been increasingly applied in hydrology to identify patterns in groundwater and streamflow time series and link them to ecosystem processes (Yu et al., 2015; Ruhí et al., 2018). A significant daily periodicity indicates that riparian vegetation is transpiring groundwater following a diel (24-hour) cycle. We ran wavelets on the longest continuous time series for each well using the ‘WaveletComp’ package (Roesch & Schmidbauer, 2018). We then horizontally “sliced” each wavelet to extract wavelet power at the daily scale across the entire study period. Using breakpoint analysis in the ‘changepoint’ package (Killick et al., 2024), we were then able to precisely determine when power at the daily scale fluctuated, pinpointing when trees began and stopped using groundwater.

To quantify groundwater use, we then followed the White method (White et al., 1932; Loheide et al., 2005) to estimate the amount of daily evapotranspiration from groundwater (*ET_gw_*, mm/d):

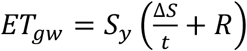

where *S_y_* is the specific yield (unitless), Δ*S* is the daily change in storage (m), *t* is the time period (24 hours), and *R* is the overnight recovery (or net inflow) rate between 0:00 to 4:00 (m/hr). The White method relies on four assumptions: (1) plant water use drives diel water table fluctuations, (2) ET is minimal between 0:00 and 4:00, (3) the rate of recharge is constant throughout both day and night, and (4) specific yield can be reliably estimated. Due to assumption 3, we did not quantify *ET_gw_* on days that it rained or the day after rain. We determined the specific yield at each well using the methods described in Loheide et al. (2005), relying on soil texture described in Scheiderich et al. (2022) and DTW. Figure 1 shows the DTW time series (Figure 1D) and representative diel water table fluctuations and the implementation of the White method (Figure 1C).

To test our second hypothesis—that depth to groundwater and precipitation are the main controls on vegetation greenness (NDVI) across flow regimes and species—we used random forest (RF) regression models. RF models have been used to identify driving forces in vegetation greenness changes (Wang et al., 2022; Qiao et al., 2024) and understand ecosystem reliance on groundwater (Ruehr et al., 2023). Using data from 2020-2024, we ran six RF models, each predicting the average NDVI for a specific flow-species group (e.g., cottonwood by perennial sites or valley oak by intermittent sites), using DTW, temperature (Temp), precipitation (Precip), one-month lagged precipitation (Precip_1m), photoperiod (Photo), and vapor pressure deficit (VPD) as predictors. For the intermittent models, DTW for the CHA study area was the average of the CHA-2, CHA-3, and CHA-4 time series. We performed a hyperparameter grid search using the ‘ranger’ package (Wright and Ziegler, 2017). We tested using 1, 2, or 3 predictors at each node split as well as a minimum node size of 1, 3, or 5, again selecting the hyperparameter combination with the lowest RMSE. The number of trees for all models was set to 500 and trained on a random 75% sample of the data. We used the ‘vivid’ package (Inglis et al., 2022) to estimate variable importance and its interaction effects for each predictor.

To test our third hypothesis that drought shortens the growing season by accelerating leaf senescence, with greater effects at intermittent sites, we compared NDVI time series across dry (2020-2022) and wet (2019, 2023, 2024) years, stratified by flow regime and species. To quantify the overall impact of drought on vegetation greenness for each flow regime-species grouping, we used Student’s t-tests to compare mean NDVI during the growing season between drought and non-drought years. Shifts in SOS, EOS, and GSL due to drought were calculated as the difference in phenophase timing between drought and non-drought years. We then sought to place our findings in a broader ecological context, comparing our results to global observations of drought impacts on phenology sourced from long-term satellite observations from the Global Inventory Modeling and Mapping Studies (GIMMS) NDVI 3g dataset (1982-2015); ground-based observational data from the Pan European Phenology Network (PEP725) (1945-2016), the Russian “Chronicles of Nature” Network (RCNN) (1901-2017), and the China Phenological Observation Network (CPON) (1963-2014) (Liu et al., 2025); and experimental precipitation manipulations (Lu et al., 2023). We used the Global Aridity Index and Potential Evapotranspiration Database (Zomer et al., 2022) to classify observations by aridity class (Humid, Dry Sub-Humid, Semi-Arid, Arid, and Hyper Arid), then ran two-way ANOVA with phenological shifts (in days) as a function of aridity classification and study type; and post-hoc Tukey tests to assess differences in phenological shifts between study types and aridity classes.

## Results

### Groundwater use by riparian trees

Annual water table fluctuations were observed at all wells (Figure 1D). Water table depths at all wells showed a clear response to precipitation patterns, with minimum depths occurring at the end of dry seasons and rising rapidly with the onset of precipitation events. The magnitude of response varied across monitoring wells, with SC-2 consistently showing the largest annual fluctuations (5.4-6.6 m), while the only perennial site SC-5 exhibited much more stable conditions (0.5-0.8 m). The CHA wells displayed intermediate fluctuation ranges (3.7-4.7 m). The interannual variation in precipitation and PDSI was strongly associated with groundwater dynamics across all monitoring sites (Figure 1B). For example, SC-4 experienced a sharp decline in groundwater depth during the 2020-2022 drought until wet conditions in 2023 recharged the aquifer.

Wavelet analysis revealed diel water table fluctuations at all wells. To illustrate the range of responses across the hydrologic gradient, we compared two wavelets—one from the well by a perennial reach (SC-5) and one from a well by an intermittent reach (CHA-4) (wavelets from the remaining intermittent wells can be found in Figure S1). At the perennial site, wavelet power at the daily period (period = 1) was the strongest from May to November every year, revealing consistent diel fluctuations during the dry season (Figure 2A). At the intermittent site, wavelet power at the daily scale was comparatively weaker but persists longer throughout the year (Figure 2B). Breakpoint analysis (Figure S2) further showed that the date at which wavelet power declined each year coincided roughly with the water table falling below ∼5m, revealing a critical threshold for groundwater access.

**Figure 2.**
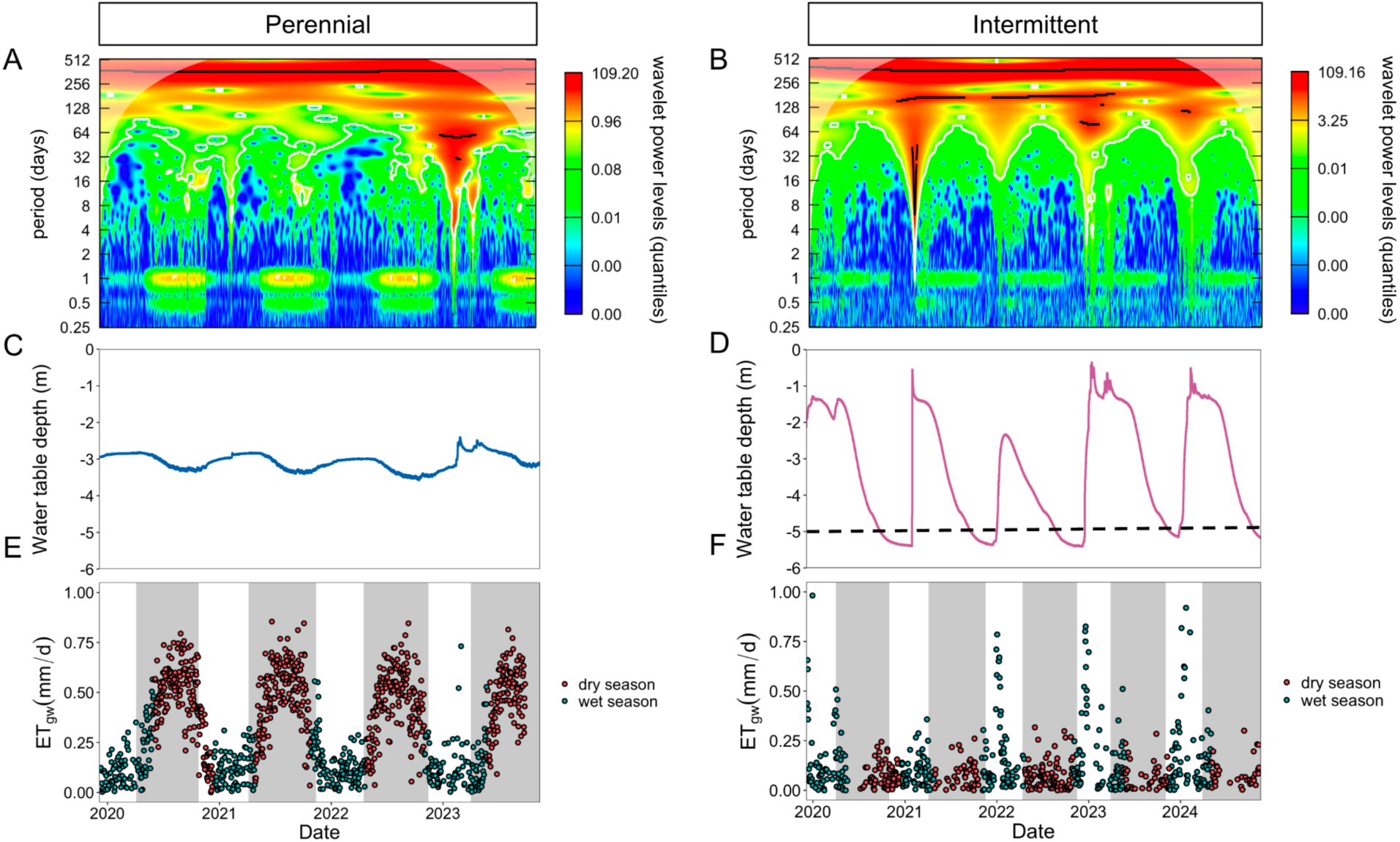
Timing and magnitude of groundwater use by trees. (A, B) Wavelet diagrams illustrating the power of each frequency in depth to groundwater data from wells SC-5 (perennial) and CHA-4 (intermittent). Warmer colors at the daily scale (period = ∼1) indicate when trees are using groundwater. (C, D) Depth to water table. The horizontal black line in panel D illustrates the estimated depth at which trees can no longer exert diel fluctuations on the water table at CHA-4. (E, F) Estimates of evapotranspiration from groundwater (ET_gw_) (mm/d) from the White method shown for the dry (red) and wet (blue) seasons. Growing seasons are highlighted in gray.

ET_gw_ also displayed a seasonal pattern at both perennial and intermittent sites. During the wet season, ET_gw_ was slightly higher at the intermittent site, averaging 0.19 mm/d compared to 0.16 mm/d at the perennial site. During the dry season, ET_gw_ increased substantially to an average of 0.49 mm/d at the perennial site while the intermittent site declined to just 0.10 mm/d—an almost fivefold difference between flow regimes (Figure 2E). These results support our hypothesis that riparian trees near perennial streams would primarily rely on groundwater during the dry summer—while trees near intermittent streams would be limited by groundwater depth, maintaining transpiration via groundwater as long as the water table was shallower than ∼5m.

### Patterns in riparian greenness and phenology

The NDVI time series for tree species within each well’s area of influence exhibited clear seasonal fluctuations, with peak values occurring during the delineated growing seasons (Figure 3A). However, the magnitude and variability of NDVI differed by tree species and site hydrology. ANOVA analyses and Tukey’s HSD post-hoc comparisons showed that the perennial site (SC-5) had significantly greater average growing season NDVI values compared to the three intermittent sites across all species (14-84% greater for willow, 6-54% greater for cottonwood, and 4-26% greater for valley oak; *p* < 0.001 for all comparisons) (Figure 3B). SC-4 consistently displayed the lowest NDVI values across all three species compared to other sites (36-46% lower for willow, 24-35% lower for cottonwood, and 11-20% lower for valley oak; *p* < 0.001 for all comparisons). These results are consistent with our prediction that perennial sites would exhibit greater and more stable greenness. Moreover, this site-based variation suggests that local hydrology and species-specific traits may modulate greenness independently of broader seasonal cues. These differences motivated our subsequent analysis of the environmental drivers shaping NDVI dynamics across sites and species.

**Figure 3.**
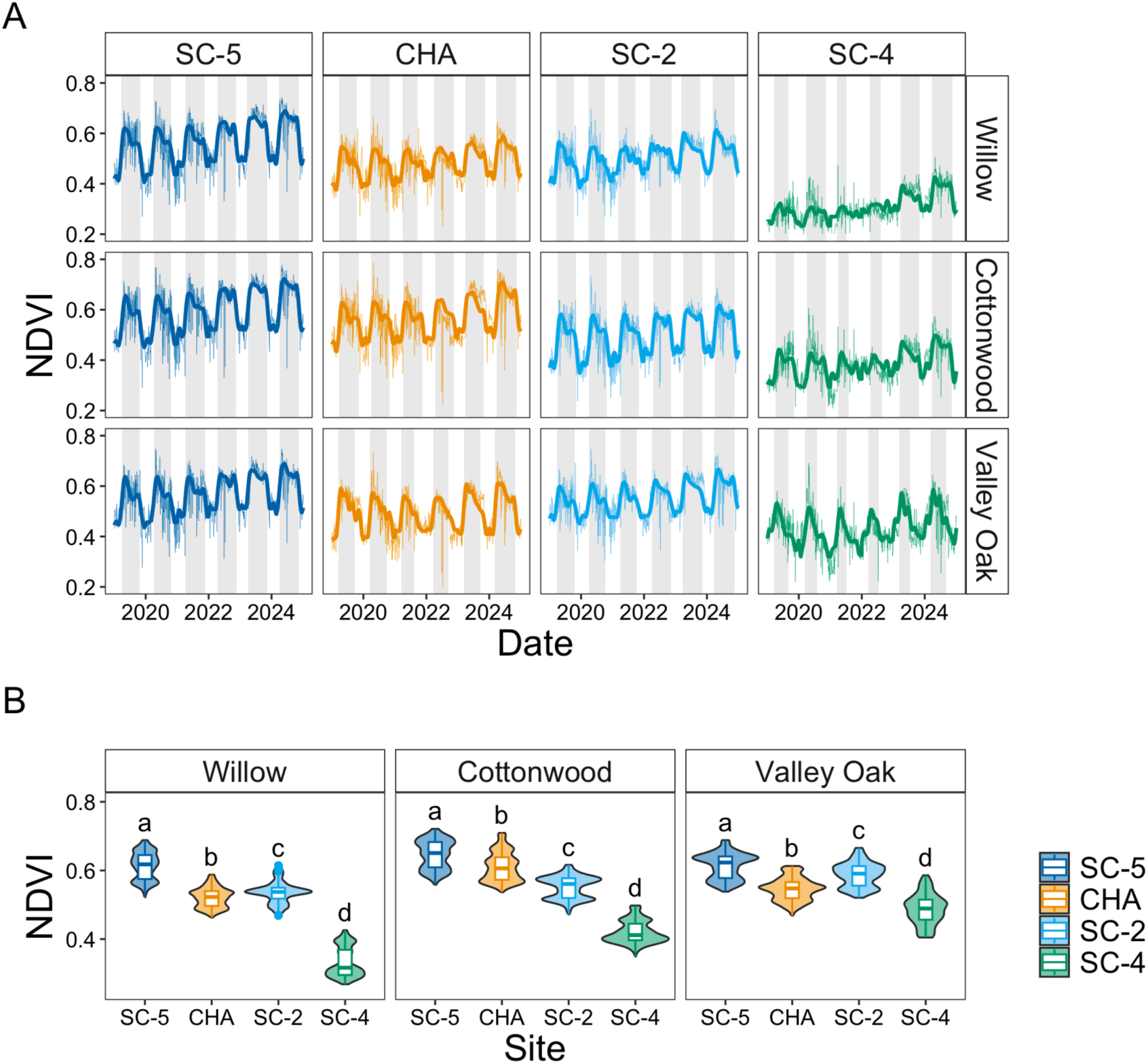
Normalized Difference Vegetation Index (NDVI) time series and growing season division from satellite imagery. (A) NDVI time series for willow, cottonwood, and valley oak trees within each site (indicated by line color). SC-5 is the only perennial site among the four, and SC-4 has the deepest groundwater. The raw average values are plotted in lighter colors with the weighted HANTS (Harmonic Analysis of NDVI Time Series) smoothed values plotted with darker, thicker lines. Growing seasons delineated using the threshold method are highlighted in gray. (B) Violin and boxplots of smoothed growing season NDVI values binned by site. Letters indicate groupings based on Tukey’s HSD test: different letters denote statistically significant differences (p < 0.05) between sites. The boxes represent the interquartile range 25th to 75th percentiles, thick lines mark the median, whiskers represent the 5th and 95th percentiles, and outlying points are plotted individually.

### Drivers of riparian greenness and phenology

The random forest models explained a high proportion of the variance (>87%) in NDVI around the wells. The primary driver of greenness differed distinctly between site types: photoperiod dominated at perennial sites, while depth to water (DTW) was the controlling factor at intermittent sites (Figure 4). At perennial sites, all three species exhibited complex interaction networks with multiple influential variables. All species showed strong interactions between photoperiod, DTW, and temperature, with photoperiod having the highest importance. Of the three species at the perennial site, valley oak had more balanced variable importance between photoperiod, PDSI and DTW. In contrast, species at intermittent sites demonstrated much simpler interaction networks dominated by DTW. Willow and cottonwood showed the strongest DTW influence and DTW-photoperiod interaction, while photoperiod and PDSI were distinctly of secondary importance compared to DTW. In contrast, valley oak at intermittent sites uniquely maintained stronger photoperiod importance alongside DTW compared to the other species, suggesting a more balanced response to both water availability and seasonal cues.

**Figure 4.**
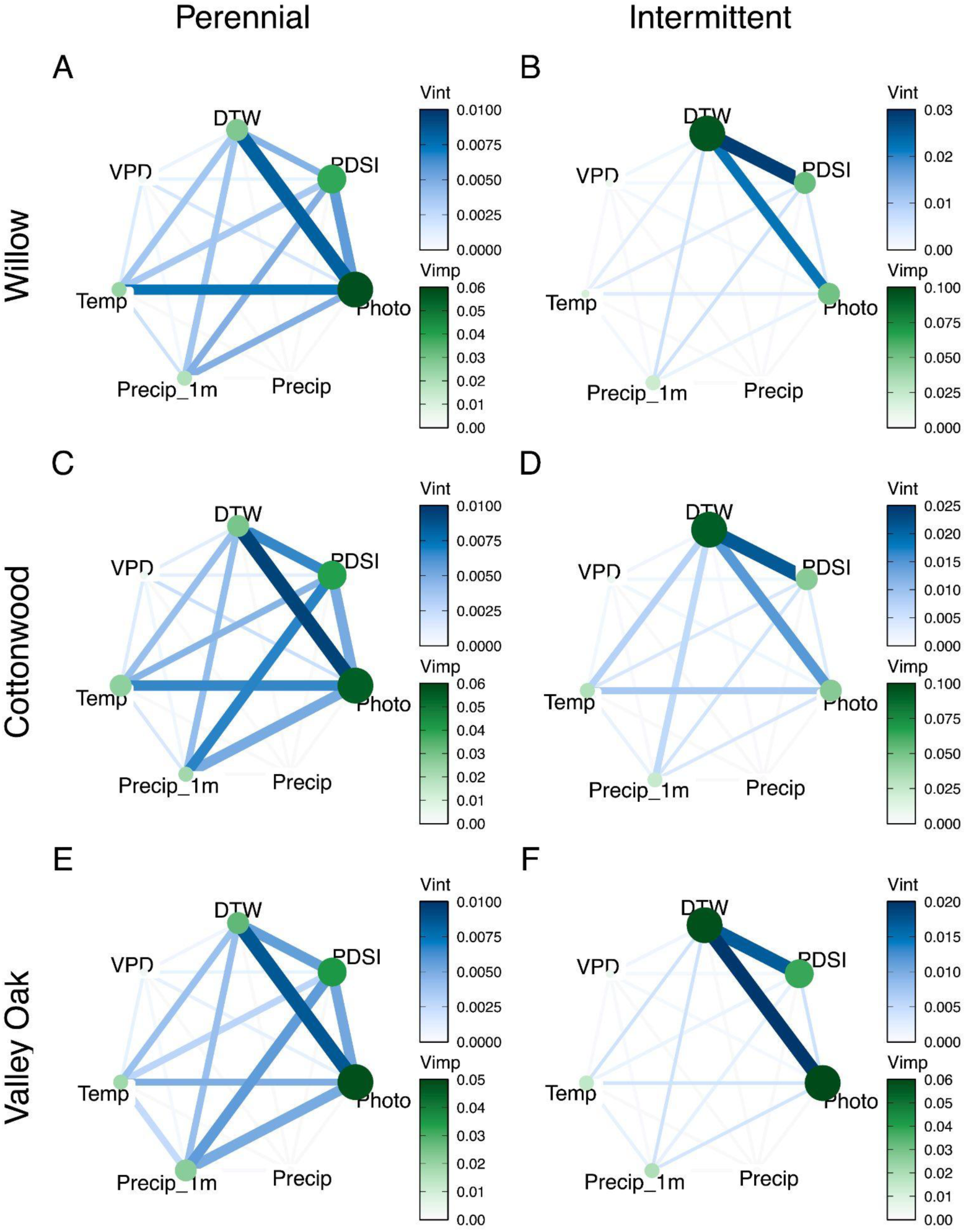
Variable importance and variable interaction network plots from random forest models predicting vegetation greenness (Normalized Difference Vegetation Index, NDVI) for perennial and intermittent cottonwood (A, B), valley oak (C, D), and willow (E, F). Higher variable importance (Vimp) is indicated with larger and darker nodes (green). Higher variable interaction strength (Vint) is plotted with thicker and darker lines (blue).

These results partially support our hypothesis regarding the importance of DTW for vegetation greenness. Specifically, while photoperiod emerged as the dominant driver of greenness at perennial sites where water availability was stable, this primary phenological control was supplanted by DTW at intermittent sites.

### Influence of drought

Historic drought conditions between 2020-2022 were associated with lower NDVI values across all species and both flow regimes (Figure 5A). Student’s t-tests showed this difference was significant for cottonwood (perennial: t_1185_ = 12.50, *p* < 0.0001; intermittent: t_3758_ = 7.53, *p* < 0.0001), willow (perennial: t_520_ = 5.52, *p* < 0.0001; intermittent: t_1232_ = 15.03, *p* < 0.0001), and valley oak (perennial: t_1222_ = 12.17, *p* < 0.0001; intermittent: t _3261_ = 13.99, *p* < 0.0001). Willow experienced the greatest NDVI declines under drought conditions (−5.18% for intermittent and - 4.78% for perennial) followed by valley oak (−4.48% for intermittent and −3.84% for perennial) and cottonwood (−3.87% for intermittent and −4.09% for perennial).

**Figure 5.**
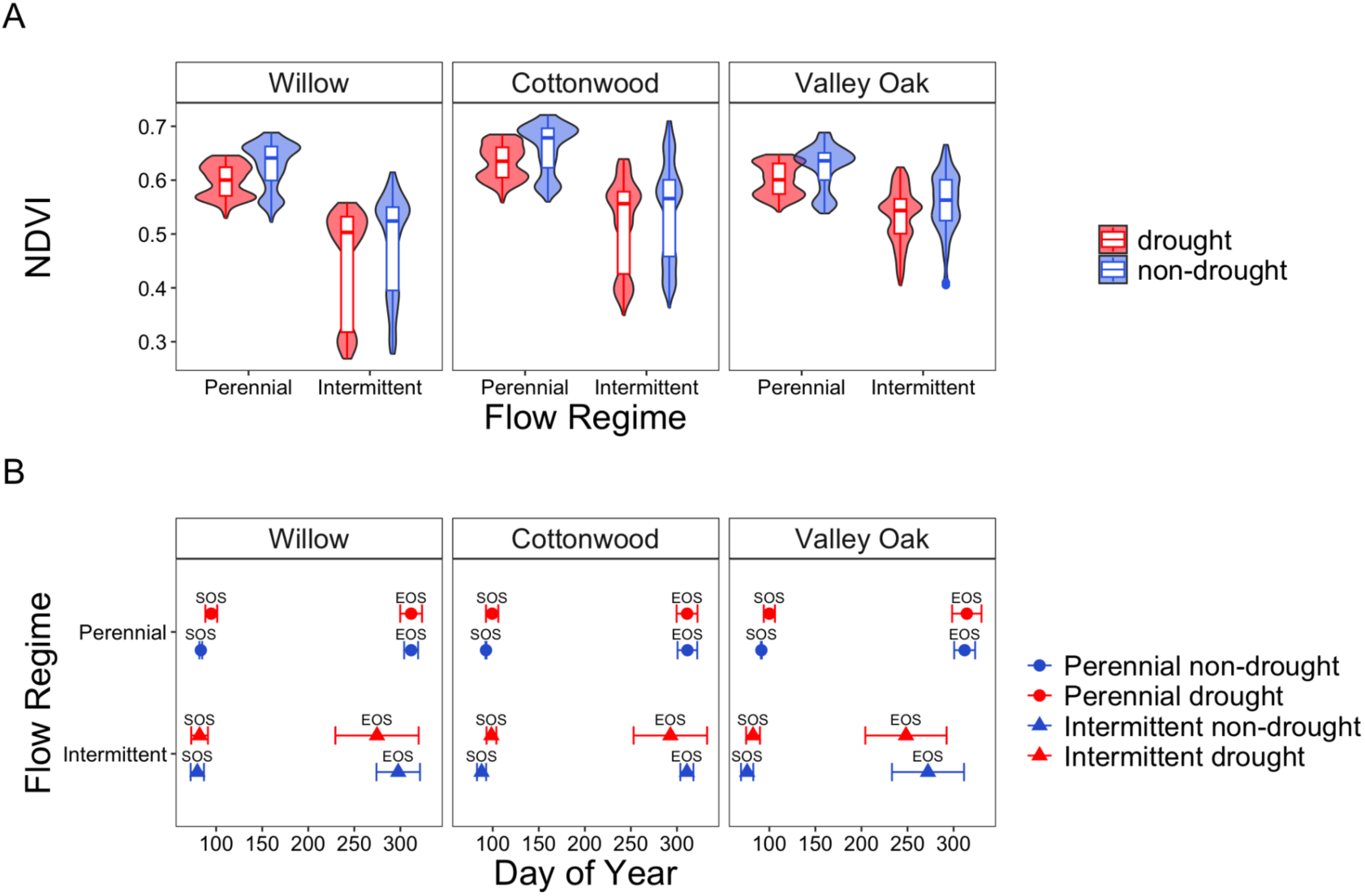
Impacts of drought on growing season Normalized Difference Vegetation Index (NDVI) and phenology metrics. (A) Violin and boxplots of smoothed growing season NDVI values (see Methods for details) for each tree species binned by flow regime and drought (2020, 2021, 2022) and non-drought (2019, 2023, 2024) years. Boxes represent the interquartile range 25th to 75th percentiles, thick lines mark the median, whiskers represent the 5th and 95th percentiles, and outlying points are plotted individually. (B) Average growing start of season (SOS) and end of season (EOS) for each tree species binned by flow regime and drought (2020, 2021, 2022) and non-drought (2019, 2023, 2024) years. Dots represent the mean and whiskers are +/- one standard deviation from the mean.

Our analysis also revealed phenological shifts across all three riparian tree species in response to drought, with notable differences between perennial and intermittent flow regimes (Figure 4B). At perennial reaches, all three species delayed their start of the growing season (SOS) during drought conditions compared to non-drought periods. Cottonwood, willow, and valley oak showed an average delay of 6.70, 11.40, and 8.73 days, respectively. End of the growing season (EOS) timing remained relatively stable at the perennial site, with minimal changes between drought and non-drought conditions (on average, willow and valley oak EOS was delayed by 0.08, and 2.55 days, respectively, while cottonwood EOS advanced by an average of 0.44 days).

Notably, trees by intermittent sites displayed more pronounced responses to drought. While SOS was also delayed during drought (by 10.65, 1.99, and 6.90 days for cottonwood, willow, and valley oak, respectively), the most dramatic changes occurred in EOS timing. All three species in intermittent flow regimes showed substantially earlier EOS during drought conditions: cottonwood by 10.85 days, willow by 15.60 days, and valley oak by 20.96 days. These phenological shifts resulted in significant changes to growing season length. In perennial systems, the growing season was shortened during drought by 7.14, 11.32, and 6.18 days for cottonwood, willow, and valley oak, respectively. The effect was more pronounced in intermittent systems, where the growing season contracted by 21.49, 17.59, and 27.86 days for the same species.

Drought-induced phenological shifts documented in this study, particularly at intermittent sites, tended to be greater than those observed in other ecosystems (Figure 6). All ecosystems experienced a delay in SOS, with differences across study type-aridity classes (F_[15,794279]_ = 92.36, *p* < 0.0001). However, the delayed SOS at intermittent and perennial sites were not significantly different compared to other ecosystems. EOS shifts showed significant variation across study type-aridity classes (F_[5,172]_ = 7.926, *p* < 0.0001), with intermittent sites experiencing the most substantial delays (*p* < 0.05 for all comparisons). Similarly, growing season length (GSL) changes differed significantly across study type-aridity classes (F_[4,69]_ = 13.21, *p* < 0.0001), with intermittent sites again exhibiting the most dramatic reductions (*p* < 0.05 for all comparisons).

**Figure 6.**
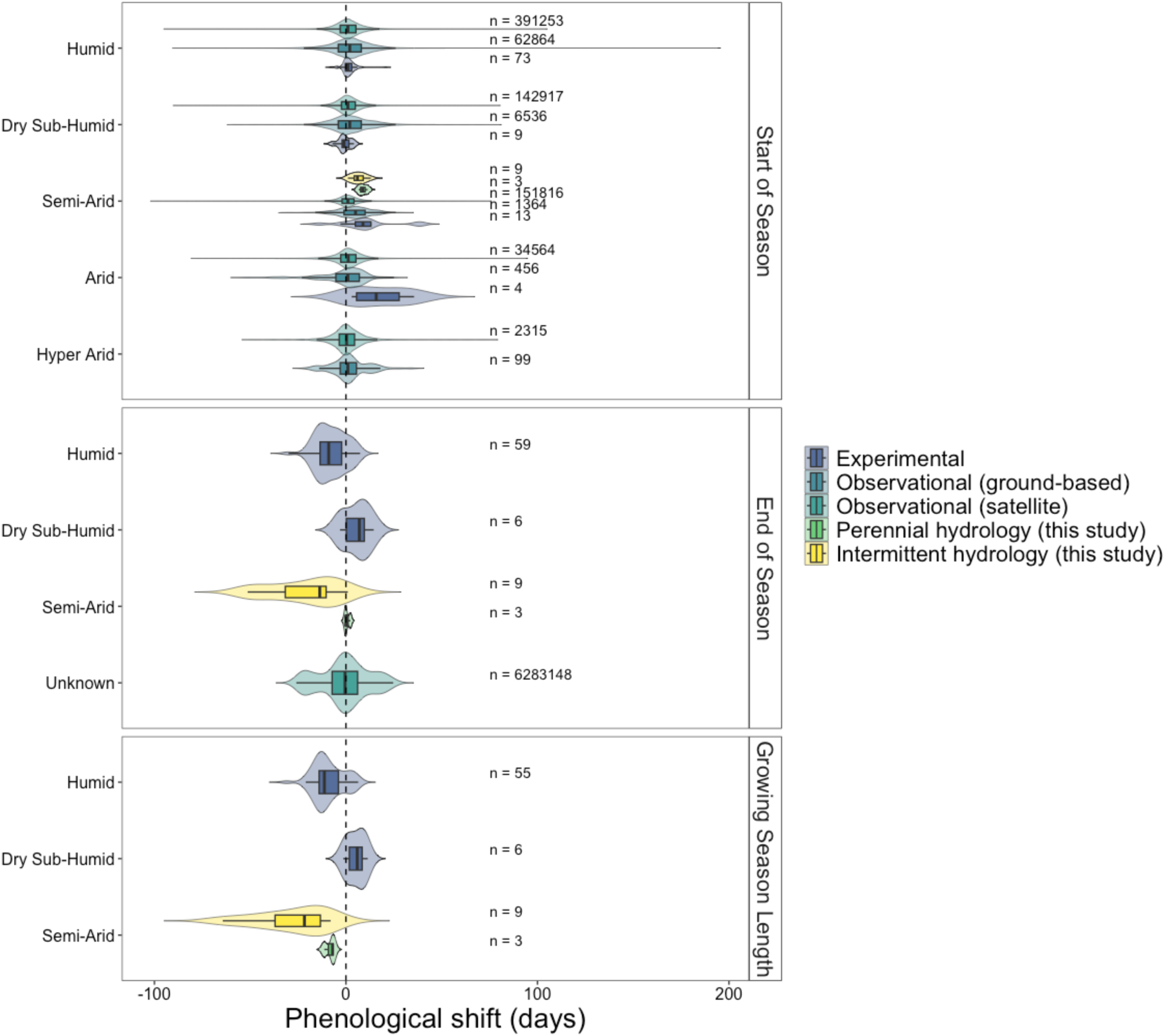
Comparison of drought-induced phenological shifts across study types and ecosystems. We compared our values on tree phenological shifts in response to drought to a large number of observation and experimental values from the literature (N= 7,077,557). We represent box and violin plots of changes in start of season (A), end of season (B), and growing season length (C), expressed as shifts in days. Colors represent different datasets: intermittent (yellow) and perennial (green) vegetation from this study; long-term satellite observations from the Global Inventory Modeling and Mapping Studies (GIMMS) NDVI 3g dataset (1982-2015) (teal; Liu et al., 2025); ground-based observational data (blue), including the Pan European Phenology Network (PEP725) (1945-2016), the Russian “Chronicles of Nature” Network (RCNN) (1901-2017), and the China Phenological Observation Network (CPON) (1963-2014), as compiled in Liu et al. (2025); and experimental precipitation manipulations (dark blue; Lu et al., 2023). Boxplots represent interquartile ranges, medians, and 5th - 95th percentiles. The dashed vertical line marks a zero shift (i.e., no change). Sample sizes (n) indicate the number of observations per group.

These findings strongly support our hypothesis that drought shortens the growing season primarily by accelerating leaf senescence, with this effect being substantially more pronounced at intermittent sites compared to perennial ones. The magnitude of phenological shifts (up to ∼28 days shortened growing season for valley oak at intermittent sites) is consistent with our prediction that dryland riparian ecosystems would be particularly sensitive to drought conditions due to their heavy reliance on groundwater, which fluctuated widely with drought intensity and duration.

## Discussion

This study examined how riparian trees in perennial and intermittent reaches use groundwater, maintain greenness, and alter their phenology in response to drought. Using wavelet analysis on long-term, high-frequency data, we found that trees near intermittent reaches experience inconsistent access to groundwater, likely reflecting the limits of their rooting depth. In contrast, trees near perennial reaches relied on groundwater primarily during the dry season, when vadose zone moisture is less available. Machine learning models on NDVI series and its potential drivers revealed that photoperiod is the dominant greenness driver at perennial sites, whereas DTW subsumes photoperiod as the primary control on greenness at intermittent ones. Drought conditions significantly reduced NDVI and growing season length across all sites, with intermittent reaches showing the most extreme phenological shifts—up to 28 days shorter. These results suggest that riparian trees by intermittent streams may be more vulnerable to water stress than previously assumed. Given the widespread, documented shifts in river flow regimes (from perennial to intermittent) across the U.S. Southwest and globally (Carlson et al. 2024), our results illustrate the significance of understanding the limits of riparian resilience to multi-year droughts.

### Riparian trees alternate between water sources seasonally

Our results suggest that trees at intermittent and perennial reaches exhibit different seasonal patterns in groundwater use. At perennial sites, the strong diel signal in groundwater levels from May to November indicates substantial groundwater extraction during the dry season, followed by a significant reduction during the wet season. This seasonal shift likely reflects changes in ecosystem water demand (e.g., leaf phenology) and the relative accessibility of shallow soil moisture and groundwater. Conversely, at intermittent sites, we observed the opposite seasonal pattern—lower groundwater use during the dry season and equivalent use to perennial sites during the wet season, corresponding to seasonal fluctuations in groundwater accessibility. Trees at intermittent sites appear to be physiologically constrained by groundwater accessibility. The apparent threshold at ∼5m water table depth beyond which wavelet power declines suggests that during the end of the dry season, the water table drops below tree’s maximum rooting depths. A similar threshold has been reported in other studies of riparian ecosystems in California (e.g., 4 m in Kibler et al., 2021; Williams et al., 2022; Rohde et al., 2024).

The Mediterranean climate context of our study area adds complexity to interpreting these patterns. Unlike temperate deciduous systems where leaf senescence coincides with increasing precipitation, this study’s riparian deciduous trees lose their leaves during the wet season and leaf out as precipitation decreases. Therefore, the period of highest transpiration demand coincides with the period of lowest precipitation input, making the relationship between groundwater use and seasonal water availability particularly important. A diel signal in DTW during the wet, leaf-off season likely represent evergreen (here, coast live oak and gray pine) groundwater use.

However, at perennial sites, possible evidence of source switching appears during the SOS. As shown in Figure 5E, during the early spring when deciduous trees resume physiological activity and begin producing leaves (increasing NDVI), ET_gw_ remains low. This suggests that despite renewed transpiration demand, trees are not yet relying on groundwater but instead utilizing available soil moisture from winter precipitation. As the growing season progresses and upper soil layers dry out, ET_gw_ increases indicating a switch to groundwater dependency. This source mixing or switching has been documented in other riparian systems (Smith et al., 1991).

The differential groundwater use of trees by perennial and intermittent reaches has profound implications for understanding climate change impacts on riparian ecosystems as previously perennial reaches may transition to intermittent flow (Zipper et al., 2021; Ayers et al., 2024; Carlson et al., 2024). Our findings suggest this shift would transform the seasonal water-use patterns of established riparian vegetation. Specifically, trees that evolved strategies optimized for perennial conditions (high dry-season groundwater use) would face significant physiological stress when forced to adopt intermittent-type strategies (reduced dry-season groundwater access), potentially leading to drought-induced mortality.

### Flow regime mediates drivers of riparian tree phenology

Our machine-learning, random forest models identified DTW, PDSI, and photoperiod as the primary drivers of riparian vegetation greenness across all studied species, though the importance of these three factors varied notably between perennial and intermittent flow regimes. At perennial sites, photoperiod was the dominant predictor of greenness whereby phenological patterns primarily follow seasonal light cues. Photoperiod is a well-documented driver of plant phenology (Piao et al., 2019) and has been observed to regulate dryland riparian vegetation phenology (McMahon et al., 2024). Despite the importance of photoperiod, the complex variable importance interaction networks observed at perennial sites indicate that these trees still respond to a suite of environmental variables including PDSI, DTW, temperature, and accumulated precipitation. In contrast, at intermittent sites, DTW largely controlled vegetation greenness, with strong interactions with PDSI and photoperiod. Recent work by Lochin et al. (2024) similarly found that late-season precipitation and groundwater availability were key to maintaining riparian greenness, and the importance of DTW in our models aligns with widespread observations of the dependence of riparian vegetation on groundwater (Stromberg et al., 1996; Pettit & Froend, 2018; Rohde et al., 2021; Williams et al., 2022). Moreover, the importance of DTW depended on species’ groundwater reliance, with anisohydric phreatophytes like willow and cottonwood showing the greatest sensitivity to DTW, and valley oak showing a more balanced response to both water availability and seasonal cues, reflecting greater drought tolerance.

A particularly strong interaction emerged between DTW and PDSI, which represent different physical and temporal scales of water availability that influence riparian vegetation in complementary ways. DTW primarily captures seasonal groundwater fluctuations linked to annual hydrological cycles and directly reflects the immediate physical constraint on root water access. PDSI incorporates longer-term moisture conditions and represents interannual climate variability that influences overall ecosystem water status. In particular, PDSI can correlate with deep (0.9-1 m) soil moisture content (Wang et al., 2015). Together, DTW and PDSI, both of which are influenced by winter precipitation, reflect root-zone soil moisture conditions. Thus, the interaction between these variables suggests that riparian vegetation responds both to immediate groundwater access as well as to unsaturated zone soil moisture. While previous studies have found that tree growth in dryland riparian systems is heavily influenced by local hydrology (Sargeant & Singer, 2021; Williams et al., 2022), our findings highlight the importance of hydrological sensitivity at multiple scales.

### Drought lowers greenness and leads to an earlier end of the growing season

The drop in NDVI values across all species during drought (2020-2022) demonstrates the vulnerability of riparian ecosystems to extended dry periods. Willow showed the most pronounced decline in greenness, followed by valley oak and cottonwood. However, these declines in greenness were relatively moderate (ranging from −3.84 to −5.18%). More revealing than absolute greenness changes were the shifts in phenological timing. Though all species showed delayed SOS during drought, the most dramatic effect was observed in EOS timing at intermittent sites, where drought triggered substantially earlier leaf senescence: 10.8 days earlier for cottonwood, 15.6 days for willow, and 21.0 days for valley oak. This asymmetric phenological response—delayed spring leaf-out but accelerated autumn senescence—resulted in significantly shortened growing seasons, particularly at intermittent sites.

The moderate greenness declines paired with the shortened growing seasons during drought suggests that at intermittent sites, trees minimized their water loss during the most water-stressed period of the year by shedding leaves, thus shifting their EOS earlier (Nolan et al., 2017; Wu et al., 2022). This plastic phenological response reflects isohydric behavior whereby trees shed their leaves to maintain more stable leaf water potentials and avoid hydraulic failure and mortality (Rood et al., 2000), especially in areas where DTW declined dramatically. For example, while all sites experienced an earlier EOS, this trend was most pronounced around well SC-4 where DTW fell nearly 5m down to 11.56m during the multi-year drought. This DTW far exceeds published maximum rooting depths reported for cottonwood (0.8-2.1m; Shafroth et al., 2000; Stromberg et al., 2013), willow (1+m; Stover et al., 2008), and valley oak (8.66m; Rohde et al., 2024).

In contrast, at the perennial site, where groundwater experienced minimal seasonal fluctuations and never declined below 3.5m, trees may have relatively constant access to groundwater to satisfy their transpiration demands despite limited precipitation and soil moisture. In this case, shallow groundwater buffered riparian vegetation from the most severe water stress (Naumburg et al., 2005; Warter et al., 2023). These trees may have employed a more anisohydric strategy, tolerating drought and maintaining stomatal conductance despite declining water availability, as their NDVI declined during drought. Still, plasticity may come at a cost. By ending their growing season earlier, trees limit carbon assimilation, potentially reducing growth and leading to carbon starvation. While this drought-avoidance strategy may help prevent immediate hydraulic damage, it may also reduce long-term fitness. Growing-season NDVI offers a proxy for photosynthetic activity, and, by extension, potential carbon gain. NDVI declines across all species and sites suggest that both tolerant (possibly anisohydric) and avoidant (isohydric) responses carry costs. Yet, the burden appears greater for trees at intermittent sites, which not only lose greenness but also growing days. This observation underscores a central trade-off: plasticity allows survival under stress but may do so at the expense of long-term productivity and resilience (Valladares et al., 2007; Vázquez et al., 2017).

## Conclusion

Riparian trees in dryland intermittent stream ecosystems have been shaped by exposure to drought over evolutionary timescales and possess adaptations to persist during periods of low water availability (Lytle & Poff, 2004; McDowell et al., 2008; Matesanz et al., 2010). However, our findings challenge the prevailing assumption that riparian ecosystems regularly exposed to hydrological stress are more resilient to drought (Munson et al., 2021; Moran et al., 2023). Instead, we found that riparian trees by intermittent stream reaches may be relatively more vulnerable to drought, curtailing the growing season via precocious leaf senescence. While drought deciduousness is a drought avoidance strategy for some dryland tree species (Dahlin et al., 2017), in our study, this premature leaf senescence at intermittent sites appears to be more of a detrimental response rather than an adaptive strategy (Moran et al., 2023). These differing responses between flow regimes are likely due to groundwater access, as trees by intermittent sites are unable to reach groundwater when the water table declines beyond ∼5m while trees at perennial sites benefit from a relatively shallow and stable water table even during dry summers (Kibler et al., 2021; Williams et al., 2022). This heightened sensitivity also suggests that riparian woodlands by intermittent rivers may be operating very close to critical groundwater thresholds, making them particularly susceptible to hydrological change. With intermittent streams comprising approximately 60% of river miles globally (Messager et al., 2021, and approximately over 81% in the arid and semi-arid American Southwest (Levick et al., 2008), vast areas of riparian woodland habitat may face under-appreciated risk (Moran et al., 2023). As non-perennial conditions are increasing in prevalence due to climate change (Zipper et al., 2021; Ayers et al., 2024; Carlson et al., 2024), trees adapted to stable groundwater conditions may struggle to acclimate, increasing the risk of physiological stress and mortality (Valladares et al., 2007). Our results highlight the need to consider flow regime shifts in predicting dryland riparian woodland resilience and underscore the importance of groundwater availability in shaping ecosystem responses to drought.

## Acknowledgements

This study was supported by a National Science Foundation (NSF) CAREER award to AR (DEB-2047324). RMM received funding from the NSF Graduate Research Fellowship Program. This study was conducted on the ancestral lands of the Amah Mutsun and Chalon peoples. We thank the staff of Pinnacles National Park, particularly Wildlife Biologist Paul G. Johnson.

