## Supplemental Figures for "Groundwater and phenology data reveal vulnerability of riparian trees to drought"

**Supporting Information**

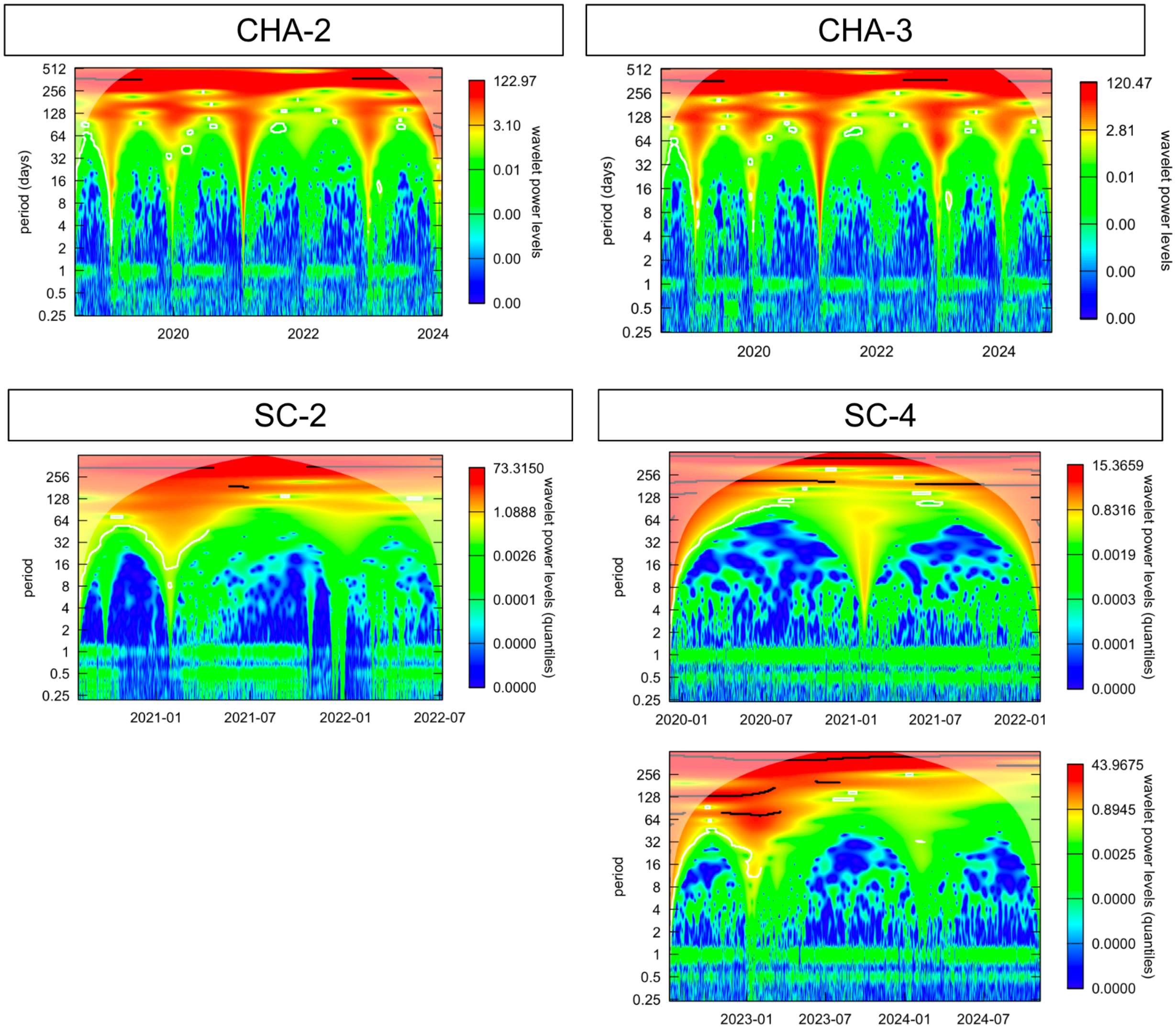


Figure S1. Wavelet diagrams illustrating the power of each frequency in depth to groundwater data from wells CHA-2, CHA-3, SC-2, and SC-4 (intermittent). Warmer colors at the daily scale indicate when trees are using groundwater.

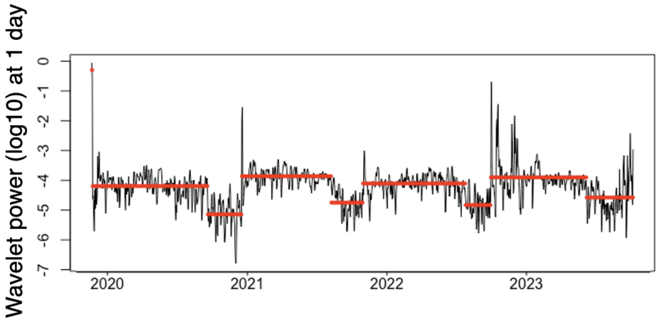


Figure S2. Daily wavelet power ($log_{10}$ scale) at the 1-day period from CHA-4. The black line shows wavelet power, while red segments indicate changepoints and mean wavelet power within each segment, highlighting shifts in diel signal strength associated with vegetation water use.
